# SpaReg: sparsity-based 3D reconstruction of tissue microenvironments at native resolution across morphological and spatial molecular modalities

**DOI:** 10.64898/2026.09.28.754175

**Authors:** Rajdeep Pawar, Thomas Jacob, Rebecca Raphael, Elaine Byrnes, Simon Watkins, T. Rinda Soong, Aatur Singhi, Shikhar Uttam

## Abstract

Tissue microenvironments comprise cellular and acellular components whose three-dimensional (3D) architecture guides disease fate. Direct imaging of intact specimens by lightsheet and multiphoton microscopy, and computational reconstruction from serial sections, have established that 3D spatial context reveals cell and tissue organization inaccessible at single planes. Computational reconstruction in particular can leverage archived human tissue, benefiting from the cost-effectiveness, robustness, scalable storage, workflow compatibility, and century-long pathobiology knowledge of histology, and can integrate multiple spatial modalities. However, sectioning can introduce tears and folds, and computational alignment can further distort tissue integrity. Here we introduce SpaReg, a sparsity-based 3D reconstruction method spanning histology, spatial proteomics and spatial transcriptomics. Across multiple organs, SpaReg robustly reconstructs large tissue volumes with preserved subcellular morphology despite sectioning artifacts. On a standardized histology benchmark, SpaReg achieves the best balance between 3D reconstruction accuracy and tissue integrity, and on spatial transcriptomics benchmarks it ranks among the leading methods while scaling to hundreds of sections and millions of cells in a dataset that several existing methods fail to process. Preservation of subcellular morphology by SpaReg also enables training of a Hematoxylin and Eosin (H&E)-based epithelial, T and B cell classifier, generating single-cell-resolved 3D maps directly from H&E. Applied to pancreatic tissue containing pancreatic ductal adenocarcinoma arising from an intraductal papillary mucinous neoplasm, these maps reveal that 2D sections overestimate immune exclusion, and resolve lymphoid aggregates in 3D. SpaReg, therefore, provides a scalable foundation for morphologically faithful, multi-modal 3D atlases and spatially informed disease modeling.

---

Understanding the spatial architecture of tissue microenvironments in three dimensions (3D) is crucial for decoding the interplay between cellular and acellular components that underlies both normal physiology and disease progression. Spatial gradients in morphology, gene expression, and cellular composition are now known to govern critical biological phenomena, including tumor evolution, immune surveillance, and therapeutic resistance (1, 2). High-resolution 3D spatial atlases have shown how phenotypically distinct but spatially contiguous structures shape tumor-immune interactions in cancer (3), exposing the limitations of conventional 2D histology in capturing graded transitions and architectural continuities. In a similar vein, 3D reconstructions of tumors across multiple types of cancer have revealed spatial subclones and microregions defined by distinct oncogenic alterations and immune niches that are poorly resolved in single-plane views (4). Such measurements can be made via direct imaging of intact specimens or computational 3D reconstruction from serial sections. Direct imaging of intact specimens by light-sheet and multiphoton microscopy recovers the microenvironment without sectioning and has enabled non-destructive whole-tissue phenotyping of 3D tumors (5–7). These approaches require specialized optical instrumentation and, depending on the imaging strategy, tissue clearing and molecular labeling(8). Reconstructing 3D tissue architecture computationally, on the other hand, from serially cut sections provides the flexibility of using Hematoxylin and Eosin (H&E), highly multiplexed fluorescence, or spatial transcriptomics(3, 9–12). Highly multiplexed fluorescence adds molecular and phenotypic detail to the reconstructed tissue that is at least an order of magnitude beyond direct 3D imaging, and spatial transcriptomics extends this to genome-scale or targeted gene expressions measured *in situ*. H&Ecaptures morphology and tissue architecture without molecular specificity, but is a cost-effective and scalable option that fits existing workflows for processing archived tissue, and draws on a century-long pathobiology knowledge base. Furthermore, its lack of molecular specificity is a gap that virtual staining and H&E-based molecular inference are beginning to close (13).

Several methods have been developed to reconstruct 3D tissue from 2D whole-slide images (WSIs) of serial sections (14–16). These methods, however, typically rely on substantial downsampling of the original high-resolution WSIs to remain computationally tractable, discarding the finer architectural and subcellular features that are critical for cell classification and spatial inference in 3D. Moreover, computational alignment can distort local tissue morphology, compromising the biological fidelity of the reconstructed tissue. A further practical limitation is that current methods commonly exclude sections affected by histological artifacts. Because these artifacts are unavoidable in standard histological workflows, excluding affected sections prevents contiguous reconstruction across large tissue specimens. The resulting gaps in 3D tissue architecture can cause spatially restricted features, such as rare cell populations, small invasive foci or localized cell–cell interactions, to be missed entirely, potentially biasing the reconstruction toward more abundant and spatially extensive structures. A parallel literature addresses the alignment of spatial-omics sections, establishing correspondence from molecular feature similarity rather than from morphology (10–12, 17–21). These methods operate on sparse measurement grids rather than gigapixel morphology, are largely tailored to individual platforms, and optimize for continuity of expression rather than morphological fidelity (22). However, a single integrated reconstruction approach that supports large-scale 3D reconstruction across histological and molecular modalities, preserves subcellular morphological detail in microscopy images, and remains robust to sectioning artifacts is still lacking.

Beyond reconstructing tissue architecture, resolving biologically meaningful cell identities within the reconstructed tissue is essential for cell-type-resolved 3D analysis of tissue microenvironments. In carcinomas, for example, resolving the spatial organization of the epithelial compartment relative to infiltrating immune populations is directly informative of immune surveillance in the tumor microenvironment, revealing which immune cells associate with or are excluded from neoplastic epithelium and how lymphoid aggregates are organized around the tumor (1, 23, 24). In the specific context of pancreatic ductal adenocarcinoma (PDAC), one of the most lethal forms of cancer (25), multiplexed spatial studies have identified clinically relevant heterogeneity in epithelial-immune organization, with greater abundance of immune-excluded tumor cells associated with shorter survival (26–28) and T-cell-dominant neighborhoods enriched for T and B cells associated with longer survival (29–31). These findings, however, were based on molecularly defined cell identities measured in individual 2D sections, which do not capture spatial relationships with cells above or below the section plane. Although deep learning methods for inferring cell identities from H&E have been developed using both expert-derived annotation (32–35) and molecularly defined labels from paired multiplexed imaging (36–39), no existing approach jointly resolves epithelial, T, and B cell identities directly from H&E tissue sections in pancreatic tissue.

To meet these requirements, we developed SpaReg, a sparsity-based, scalable method for high-fidelity 3D reconstruction that combines sparse tissue-boundary correspondence across histological and molecular modalities with subsequent refinement for accurate 3D reconstruction. We also developed a SpaReg enabled H&E-based cell classification model for epithelial, T, and B cell identities trained on molecularly defined labels derived from paired multiplexed immunofluorescence in PDAC tissue. Applying this cell classification model to reconstructed pancreatic tissue with PDAC arising from an intraductal papillary mucinous neoplasm (IPMN) (40), we recover 3D tumor-immune organization across PDAC and IPMN compartments, show that 2D sections overestimate immune exclusion, and resolve the 3D spatial organization of lymphoid aggregates.

## Results

### An overview of SpaReg

SpaReg reconstructs tissue microenvironments in 3D from serial sections, bringing H&E histology, multiplexed immunofluorescence and spatial transcriptomics within a common framework. It first recovers the section sequence from slide-label information and then registers the ordered sections through sparse tissue-boundary correspondences (Fig. 1a, Methods).

**Fig. 1.**
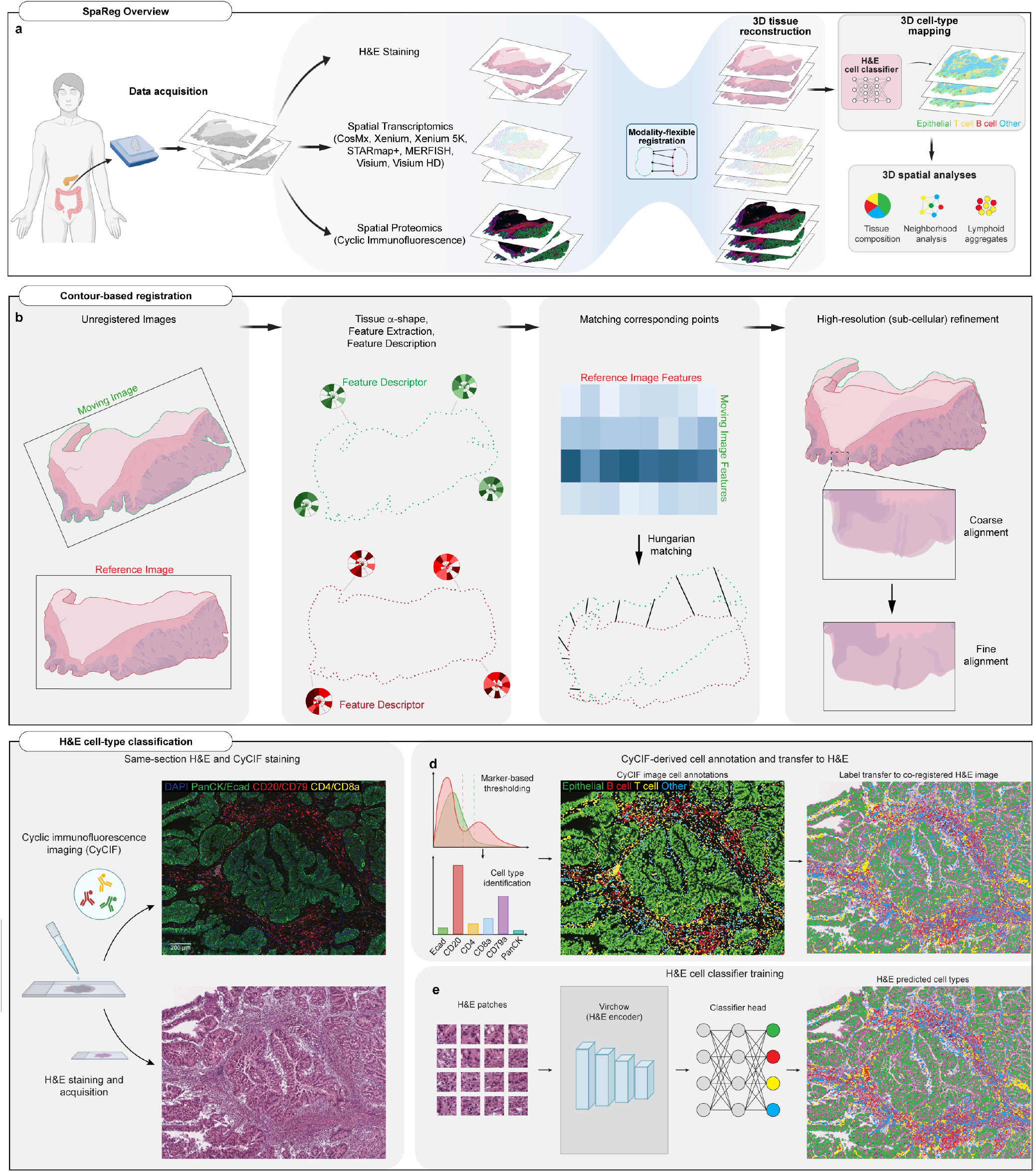
SpaReg provides a sparsity-based, multi-scale framework for 3D tissue reconstruction with cell-type-resolved microenvironment analysis. **a** Overview of 3D reconstruction from serial H&E images, spatial transcriptomics (CosMx, Xenium, Xenium 5K, STARmap+, MERFISH, Visium, Visium HD), or spatial proteomics (CyCIF) sections, along with cell-typing, and analysis of tissue composition, cellular neighborhoods, and lymphoid aggregate organization in the reconstructed tissue. **b** Tissue is represented using *α*-shape, described using shape-context features, and matched between sections by Hungarian assignment. The resulting coarse transformation identifies corresponding regions for extraction at the native resolution and subsequent local alignment refinement. **c** Paired cyclic immunofluorescence (CyCIF) and H&E imaging of the same pancreatic tissue section provides molecular reference labels for H&E-based cell classification. **d** Marker-specific thresholding assigns CyCIF-derived epithelial, T-cell, B-cell, and other-cell labels to corresponding cells in the co-registered H&E image. **e** Classifier training within the CellViT++ framework.A frozen Virchow encoder generates cell associated embeddings from H&E patches, and a multilayer perceptron predicts the four cell classes. Schematic elements were created with BioRender.com.

Two ideas separate SpaReg from existing reconstruction methods. First, initial correspondence between adjacent sections is computed from a sparse set of automatically identified tissue-boundary keypoints, derived either from down-sampled tissue images or from spot or cell coordinates in spatial transcriptomics. SpaReg extracts the tissue boundary as an *α*-shape (41), a geometric representation of the tissue outline, and represents this boundary using at most 1,000 keypoints. Matching these points between neighboring sections establishes a coarse alignment. For example, the PDAC dataset covers approximately 2.77 cm^2^ per section at an acquisition pixel size of 0.25 *µ*m/px, corresponding to approximately 4.4 *×* 10^9^ image pixels. Representing this section with 1,000 boundary keypoints gives a sparsity level of approximately 0.000023% and a data-compression factor of approximately 4.4 million for coarse alignment in this step. (Methods). Second, SpaReg reformulates registration at the native image resolution (*µ*m/pixel at which the image was acquired) as a coordinate-mapping problem rather than a whole-image warping problem. SpaReg applies the inverse transform obtained from the coarse alignment to the ROI coordinates, identifying the corresponding tissue region in the section being registered. This region is then retrieved directly from the original slide image (Fig. 1b; Extended Data Fig. 1h–l). The recovered cellular and subcellular detail is used to refine the alignment, correcting residual local misalignment without requiring whole-slide resampling. Because the ROI is specified by a fixed number of coordinates, this coordinate mapping requires only *O*(1) computation with respect to the ROI dimension. Warping an *N×N* ROI, by contrast, requires manipulating *N*^2^ pixels. Taken together, both design principles help in significantly reducing the computational burden of reconstructing 3D tissue microenvironments.

The sparse boundary representation also helps SpaReg maintain stable alignment despite localized tears, folds, debris, and non-tissue slide markings (Extended Data Figs. 1 and 2). We further demonstrate this robustness in a cholangiocarcinoma specimen comprising 80 sections, each covering approximately 5.3 cm^2^ (Supplementary Table 1). Despite substantial misalignment and several severely disrupted sections, SpaReg retains the affected sections in a coherent 3D tissue reconstruction. Supplementary Video 1 shows the original whole-slide images alongside corresponding SpaReg-registered regions across the serial sections.

The ability of SpaReg to preserve subcellular morphology makes it compatible with accurate 3D spatial analysis. To demonstrate that we developed an H&E-based classification model that identifies epithelial, T, and B cells, pooling all remaining cells into an ‘other’ category (Fig. 1c–e). For training, we performed paired cyclic immunofluorescence (Cy-CIF) and H&E imaging on the same tissue section (9, 42) to generate per-cell annotations matched to H&E (Fig. 1c–d, Methods). These labels were used to train a CellViT++ model (39) in which a frozen Virchow foundation model (43) generates per-cell embeddings and a lightweight multilayer perceptron classifies each cell from H&E morphology alone (Fig. 1e).

### SpaReg robustly reconstructs 3D microenvironments across diverse tissue types, modalities, and platforms

We first demonstrated SpaReg in a PDAC specimen reconstructed from 320 serial sections spanning approximately 25 mm *×* 18 mm *×* 1.6 mm. The H&E images were acquired at 0.25 *µ*m per pixel, and nuclear detail remained discernible throughout the reconstructed tissue (Fig. 2a, Methods). Mapping approximately 280 million classified cells revealed the 3D distribution of tumor epithelial cells and surrounding T and B cell infiltrates (Fig. 2b). A colonic specimen of colorectal cancer (CRC), reconstructed from 307 serial sections spanning approximately 28 mm*×* 16 mm *×* 1.5 mm and comprising approximately 434 million classified cells, likewise preserved tissue morphology and the spatial arrangement of these populations (Fig. 2c,d).

**Fig. 2.**
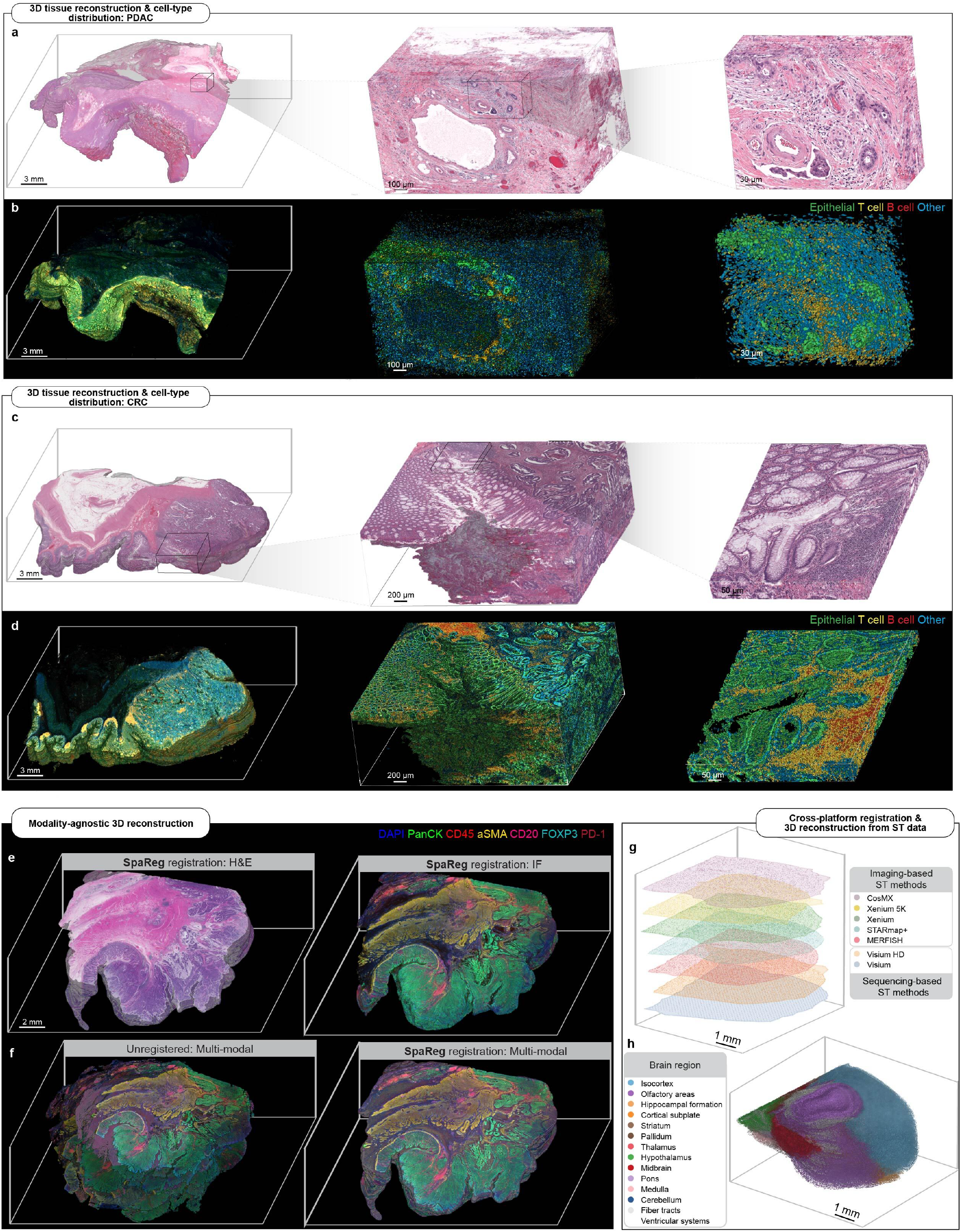
SpaReg reconstructs tissue architecture and cellular organization across tumors, modalities and platforms. **a, b** Pancreatic tissue containing pancreatic ductal adenocarcinoma (PDAC), reconstructed from 320 serial H&E sections. H&E views show glandular architecture and nuclear detail at increasing magnification (a), with corresponding H&E-predicted cell-type maps showing the distribution of epithelial cells and surrounding immune populations (b). **c, d** Colorectal tissue containing colorectal cancer (CRC), reconstructed from 307 serial H&E sections. Enlarged H&E views resolve the 3D organization of colonic crypts (c), with matched cell-type maps (d). In b and d, epithelial cells are green, T cells yellow, B cells red, and other cells blue. **e** H&E (left) and multiplexed immunofluorescence (IF, right) views of a CRC specimen from the Human Tumor Atlas Network, reconstructed from interleaved 21 H&E and 24 IF sections. Displayed IF markers are DAPI, pan-cytokeratin (PanCK), CD45, *α*-smooth muscle actin (*α*SMA), CD20, FOXP3, and PD-1. **f** Combined H&E and IF views of the interleaved sections before (left) and after (right) SpaReg based 3D registration. Corresponding 3D views are shown in Supplementary Video 3. **g** Alignment of mouse-brain sections profiled with seven spatial transcriptomics platforms, colored by platform. These comprise five imaging-based platforms (CosMx, Xenium 5K, Xenium, STARmap+, and MERFISH) and two sequencing-based platforms (Visium HD and Visium). **h** Mouse-brain hemisphere reconstructed from 129 MERFISH sections containing approximately 2.64 million cells, colored by anatomical region. Scale bars are indicated in the respective panels.

The reconstruction also recovered the native organization of colonic crypts. Although individual sections can depict crypts as separate epithelial rings, depending on section orientation, the 3D reconstruction resolved them into continuous, tube-like glands extending through the mucosa toward the muscularis mucosae, consistent with their *in vivo* anatomy. SpaReg also resolved heterogeneous tumor-stromal architecture in high-grade serous ovarian carcinoma (HG-SOC) and intricate mucosal folds in benign fallopian tube tissue (Extended Data Fig. 3a,b). Supplementary Video 2 shows successive views through the enlarged HGSC region illustrated in Extended Data Fig. 3a.

To demonstrate multimodal reconstruction, we used a lymphoid-rich specimen from the Human Tumor Atlas Network (HTAN) database (3), in which routine serial H&E sections were interleaved with highly multiplexed immunofluorescence (IF) sections resolving protein-expression-defined cell phenotypes. SpaReg registered the H&E and IF sections in their acquired interleaved order, integrating tissue architecture and cellular morphology with protein-marker distributions in 3D (Fig. 2e,f; Extended Data Fig. 3c,d; Supplementary Video 3). Cross-modal correspondence does not require matching stain appearance: each section is first aligned using shared tissue-boundary geometry, and the alignment is then refined using the common nuclear signal, the hematoxylin component for H&E and DAPI for IF.

SpaReg extends this shared geometric framework to spatial transcriptomics (ST), where differences in spatial resolution, molecular coverage and assay design make cross-section alignment particularly challenging. Sequencing-based platforms such as Visium (44) provide broad transcriptome coverage at multicellular resolution, whereas imaging-based platforms such as MERFISH (45) localize targeted transcripts at single-cell or subcellular resolution. A recent benchmark identified two persistent challenges for ST alignment: scaling ST registration to large serial-section datasets, and aligning tissue across technological platforms (22). SpaReg addressed the first by reconstructing a 129-section mouse-brain MERFISH dataset containing approximately 2.64 million cells profiled across 1,122 genes(Fig. 2h). Among the ST-specific methods we evaluated, only Spateo and SPACEL completed the registration, and both recovered section orientation substantially less accurately than SpaReg. PASTE2 and CAST, on the other hand, did not run to completion (Extended Data Fig. 4). SpaReg addressed the second challenge by bringing serial mouse-brain sections from seven spatial platforms, five imaging-based and two sequencing-based, into a shared anatomical coordinate frame (Fig. 2g). These datasets were drawn from independent published studies (46–48).

### SpaReg achieves high 3D reconstruction accuracy while preserving tissue integrity

Methods that force adjacent sections into tighter alignment can deform genuine biological differences between them, particularly when a structure is present in one section but absent from the next. In such cases, tissue may be stretched or compressed to create an artificial correspondence, altering the shape of glands, stromal regions, and tissue boundaries. SpaReg addresses this trade-off by establishing correspondence between a sparse set of keypoints sharing common structural topology across adjacent sections, while allowing genuine differences in morphology and architecture between sections to be retained (Extended Data Fig. 5).

We assessed two complementary measures: accumulated target registration error (ATRE), which captures registration drift through the serial-section stack, and tissue integrity after registration (Methods). On a standardized serial H&E benchmark, both the *α*-shape alignment (coarse) and the subsequent refinement (fine) achieved the strongest overall balance between these measures among the methods evaluated (Fig. 3a). CODA (14) introduced local distortions that altered glandular and fibroadipose architecture, whereas SpaReg preserved the original morphology while maintaining alignment (Fig. 3b). A similar contrast was observed with VALIS (15), whose global deformation visibly distorted the tissue boundary in regions that SpaReg left intact (Extended Data Fig. 6). On 10x Visium sections of human dorsolateral prefrontal cortex (DLPFC), SpaReg ranked among the top methods on both spatial-landmark accuracy and gene-expression similarity, and recovered continuous cortical layers (Layer 1 through Layer 6 and white matter) in 3D (Fig. 3c). On the same series, SpaReg limited cumulative registration drift to 0.29 normalized tissue-radius units, approximately 33-fold lower than the unaligned stack (9.60), where one tissue-radius unit is the median root-mean-square distance of spots from the centroid of each unaligned section (Methods). 3D spatial-domain clustering gave an overall score of 0.659, second to SPACEL (0.669), above PASTE2 (0.656), and substantially above the unaligned baseline (0.427) (Extended Data Fig. 7). On MERFISH sections of the mouse hypothalamic preoptic region (MHPR), SpaReg performed comparably to the leading methods and assembled the scattered input sections into a coherent anatomical reconstruction with a continuous third ventricle (V3) and fornix (fx) (Fig. 3d).

**Fig. 3.**
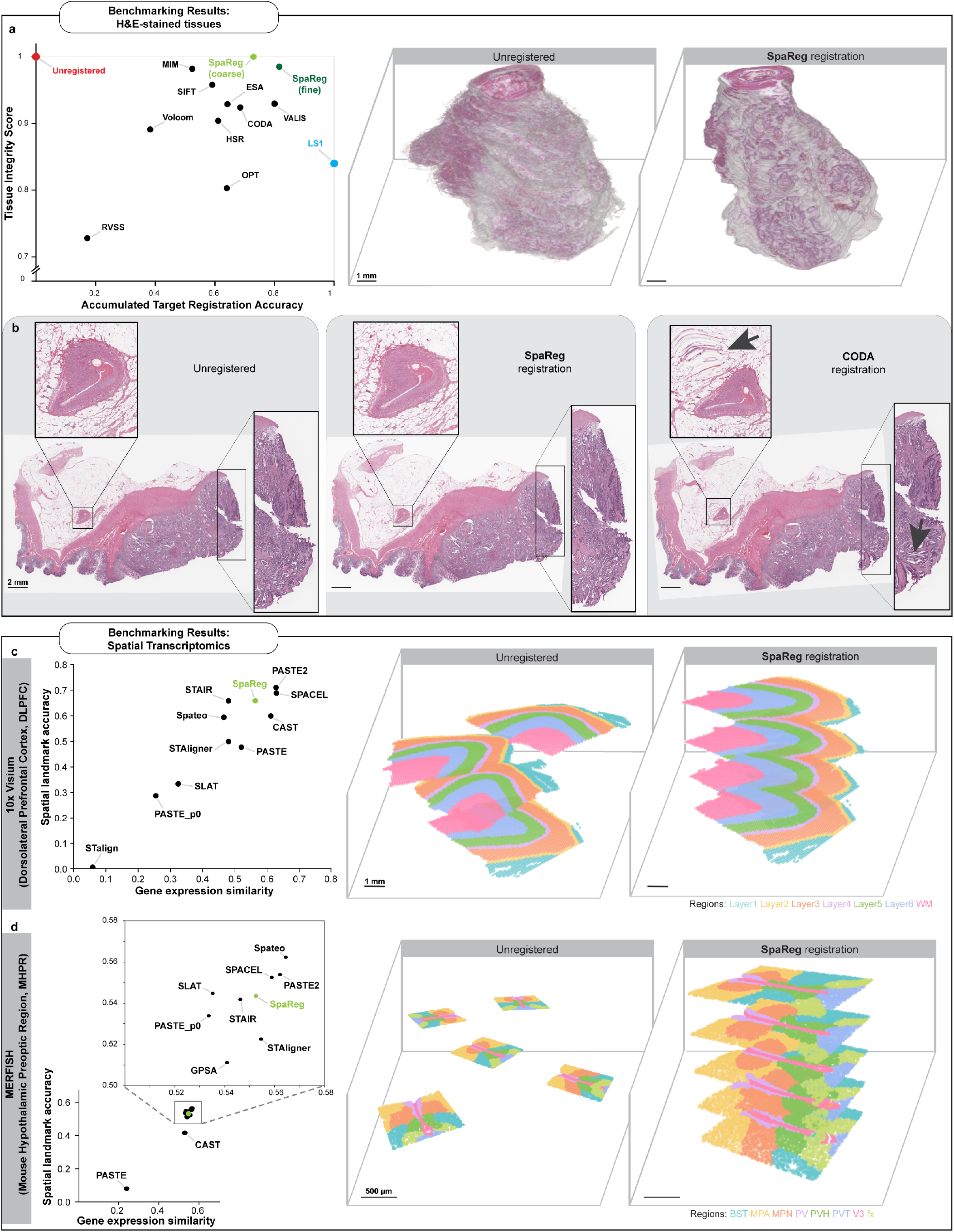
SpaReg achieves high reconstruction accuracy while preserving tissue integrity across histology and spatial transcriptomics. **a** Registration benchmark using 260 serial H&E sections of murine prostate. Accumulated target registration accuracy summarizes the reduction in cumulative landmark error relative to the unregistered and landmark-informed references. Tissue integrity score summarizes preservation of section area. Higher values indicate better performance on both axes. SpaReg is evaluated after coarse boundary registration and after image-based refinement. Representative 3D reconstructions are shown before and after registration. **b** Matched views of an original H&E section and its SpaReg and CODA registrations. Enlarged regions show glandular and fibroadipose architecture, with arrows marking local deformation in the CODA result. **c, d** Spatial transcriptomics benchmarks using four 10x Visium sections of human dorsolateral prefrontal cortex (DLPFC, c) and five MERFISH sections of the mouse hypothalamic preoptic region (MHPR, d). Gene-expression similarity measures concordance of spatial expression patterns, and spatial-landmark accuracy measures agreement of annotated anatomical regions. Higher values indicate better performance on both axes. Reconstructions before and after SpaReg registration are colored by anatomical region, showing continuity of cortical layers and white matter in DLPFC and of the third ventricle and fornix in MHPR. WM, white matter. V3, third ventricle. fx, fornix. Scale bars are indicated in the respective panels.

### Preserving spatial detail supports cell-resolved analysis in 3D

Cell-resolved 3D analysis depends on both reliable cell identities and preservation of the spatial detail needed to distinguish individual cells. The H&E classifier used to generate these 3D cell maps was trained and evaluated on 181,976 unique CyCIF-labeled cells across eight regions from a pancreatic tissue containing PDAC, using four-fold region-level cross-validation in which six regions were used for training and two were held out for evaluation in each fold. Grid-based patch extraction yielded 224,657 cell-label instances (Fig. 4a, left). The classifier achieved an overall macro F1 score of 0.663 (Fig. 4a, center). Performance was highest for epithelial cells (F1 = 0.940), followed by T cells (0.661) and B cells (0.488). The confusion matrix showed that B cells were most frequently misclassified as T cells (Fig. 4a, right). Predicted cell maps in held-out regions showed strong spatial agreement with ground truth for epithelial cells and broadly preserved lymphocyte-dense regions (Fig. 4b).

**Fig. 4.**
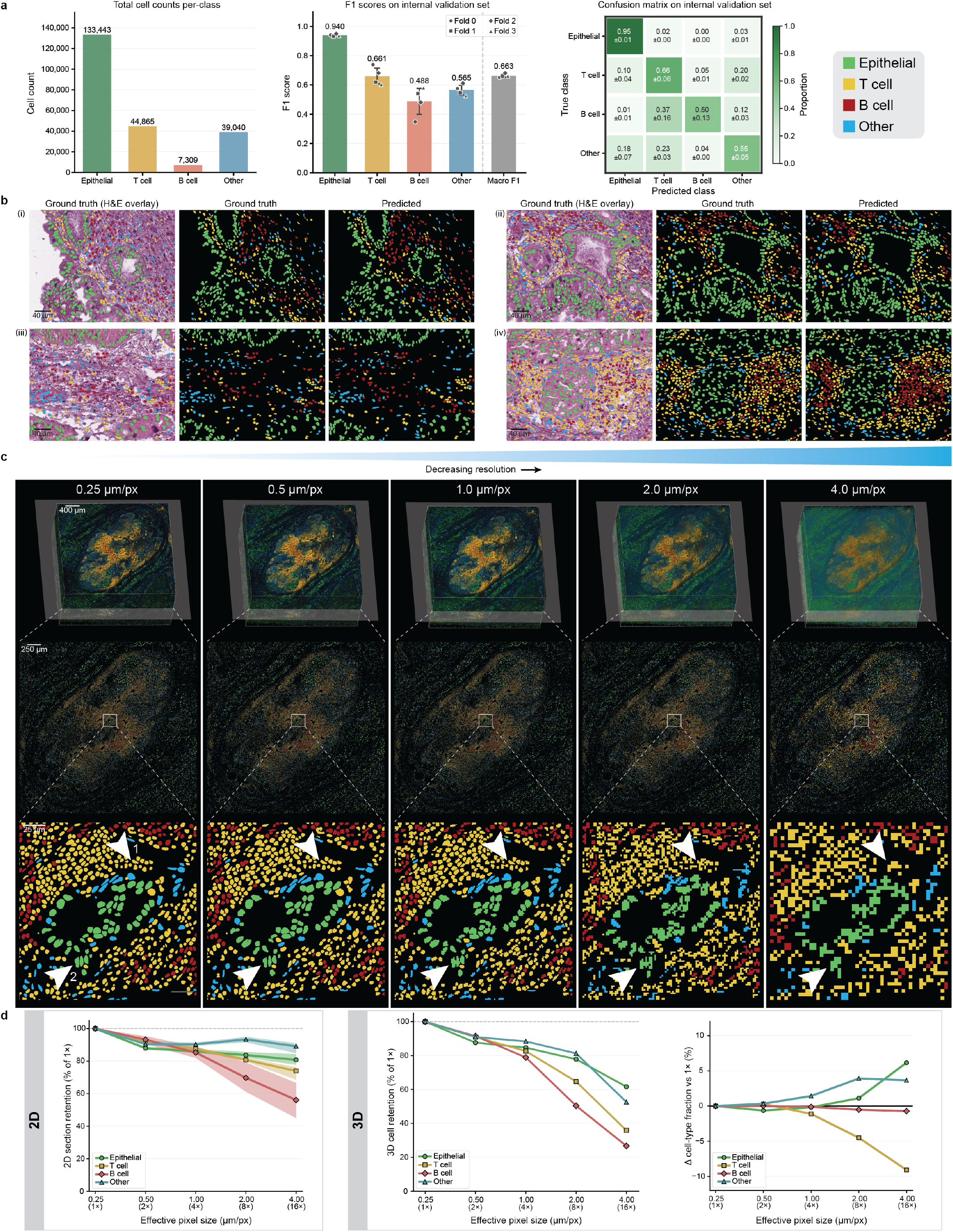
Native-resolution reconstruction preserves single-cell detail in H&E-derived 3D maps. **a** Number of cell label instances per class used for training and evaluation (left). Per-class and macro F1 values from four-fold region-level CV with bars representing mean across folds, error bars representing s.d., and individual data points per fold (center). Confusion matrix row-normalized within each fold and averaged across folds with mean *±* s.d. (right). **b** Four representative fields chosen from held-out validation regions to demonstrate classifier performance versus ground truth. **c** 3D reconstructed cell maps of PDAC-containing pancreatic tissue following progressive downsamplng of classified cell instance masks from original pixel resolution of 0.25 *µ*m/px to effective pixel resolutions of 0.5, 1.0, 2.0, and 4.0 *µ*m/px (top). Representative 2D sections from the 3D volumes (middle). Zoomed-in views of a PDAC-containing ROI from corresponding 2D sections (bottom). White arrowheads depict example sites across all chosen effective pixel resolution, with arrowhead 1 showing adjacent T cells merging into a single retained connected component with decreasing effective pixel resolution, and arrowhead 2 showing neighboring epithelial cells merging with a smaller adjacent T cell, causing the resulting component to be assigned to the epithelial class. **d** Summary of cell type counts at each pixel size as a percentage of total cell counts at 0.25 *µ*m/px across 320 serial sections, with solid lines representing the median count and shaded bands representing the IQR (left). Cell type counts at each pixel resolution across the reconstructed volume as a percentage of the counts at 0.25 *µ*m/px (center). Signed deviation of the cell type composition of the reconstructed volume at each pixel size from the original cell type composition at 0.25 *µ*m/px, reported in percentage points (right).

We next examined how coarser spatial sampling affects these cell-resolved maps. We analyzed a 15.6 mm^3^ pancreatic specimen containing PDAC spanning 1.595 mm in depth, progressively downsampling its classified cell instance masks from the original pixel size of 0.25 *µ*m to effective pixel sizes of 0.5, 1.0, 2.0, and 4.0 *µ*m and quantifying the number of distinct cells retained at each pixel size (Methods). Cell counts declined steadily with increasing effective pixel size, with reduced counts already detectable at 0.5 *µ*m/px and becoming progressively more pronounced from 1.0 *µ*m/px onward (Fig. 4c), both per section (Fig. 4d, left) and across the reconstructed volume (Fig. 4d, center). These reductions were cell-type dependent, with lymphocytes the most affected population, altering the apparent cell-type composition of the tissue (Fig. 4d, right). Visual inspection showed that closely packed cells merged into single connected components as pixel size increased, and that smaller lymphocyte profiles were more likely to merge with larger adjacent epithelial profiles, with the merged object assigned to the epithelial class (Fig. 4c, arrowheads 1 and 2). This compositional drift is consequential because the preferential loss of small lymphocytes biases cell-type-resolved measurements, including the intratumoral T cell densities that carry established prognostic consequences (1, 23).

### 3D reconstruction corrects 2D overestimation of immune exclusion and recovers known IPMN–PDAC differences from H&E alone

We compared the same tissue as independent 2D sections and as a reconstructed 3D volume to determine how sampling geometry influences biologically relevant measurements. Within a 120 mm^3^ SpaReg-reconstructed pancreatic tissue volume, we performed two complementary analyses: T cell proximity to epithelium in PDAC and T cell–epithelial organization across high-grade IPMN and IPMN-associated PDAC (Fig. 5a).

**Fig. 5.**
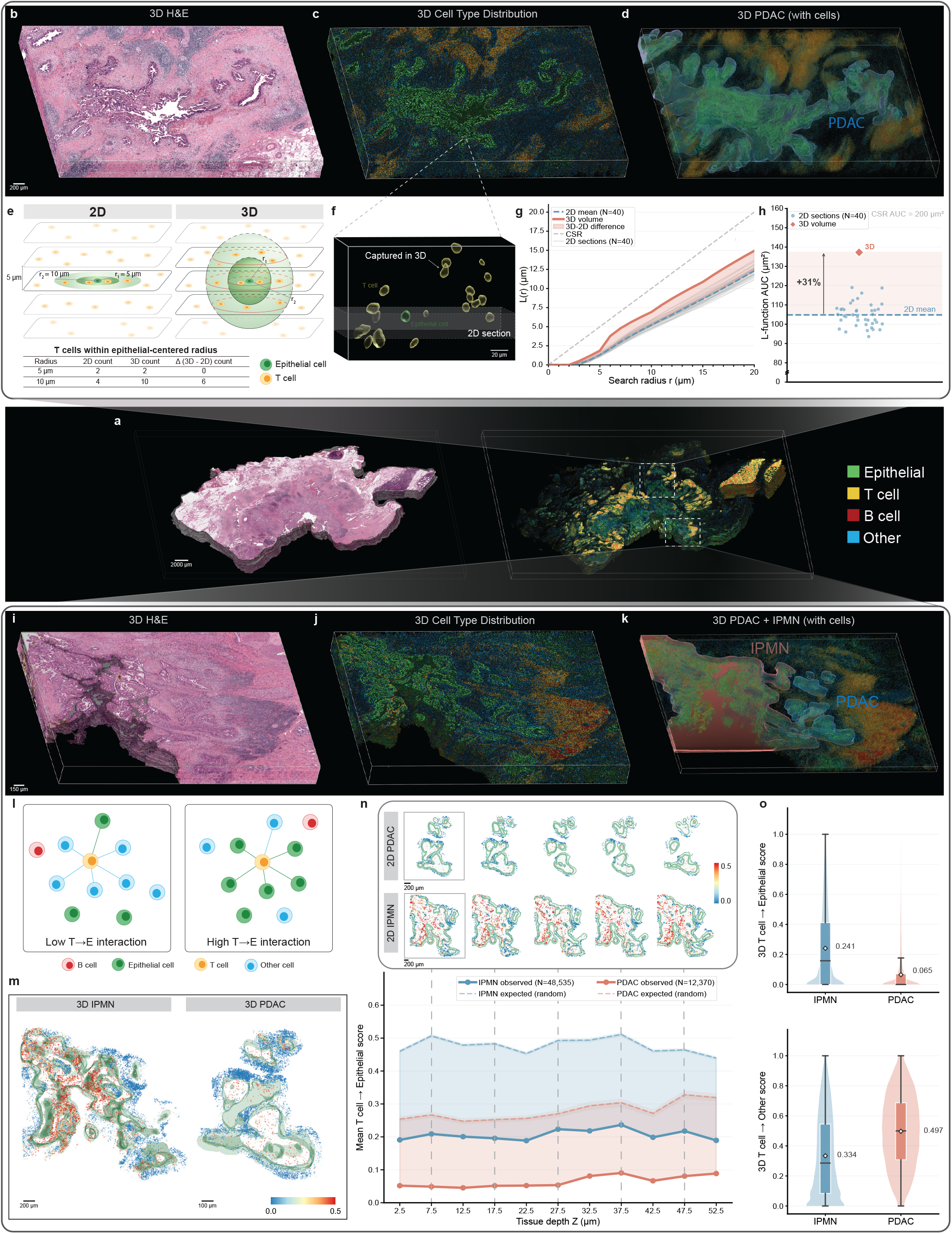
3D reconstruction reveals that 2D sections overestimate immune exclusion in PDAC and shows greater T cell exclusion in PDAC than in IPMN. **a** Pancreatic tissue reconstructed in 3D by SpaReg. H&E (left). Cell type distribution (right). **b**-**d** PDAC-containing ROI. **b** 3D H&E reconstruction. **c** 3D cell type map. **d** 3D PDAC compartment. **e** Schematic representing the 2D versus 3D cross-L function comparison. **f** Zoomed-in ROI illustrating an epithelial-centered 3D neighborhood and T cells captured above and below the corresponding 2D section. **g** Epithelial to T cell cross-L function *L*(*r*) over 0-20 *µ*m radius using epithelial cells within the PDAC annotation and T cells within the PDAC annotation and its surrounding 20 *µ*m margin. **h** L-function area under the curve (AUC) over 0-20 *µ*m. **i**-**j** ROI with PDAC and IPMN compartments. **i** 3D H&E reconstruction. **j** 3D cell type map. **k** 3D PDAC and IPMN compartments. **l** Schematic representing T cell to Epithelial (T *→*E) interaction score calculation. **m** 3D T *→*E interaction score maps for IPMN and PDAC compartments. **n** PDAC and IPMN T *→*E interaction score spatial maps for few 2D sections across tissue depth (top). Mean T *→*E interaction score per section versus tissue depth for PDAC and IPMN (bottom). Solid lines - observed within-section mean, dashed lines - random-mixing null. **o** Per T cell 3D interaction score distributions in PDAC and IPMN microenvironments. T *→*E (top) and T *→*Other (bottom). Violin plots - full per-cell distributions, white diamonds - mean, box plots - median and IQR.

Cytotoxic T cell proximity to cytokeratin 8^+^ cancer cells, quantified by the area under the curve (AUC) of Ripley’s L-function (49, 50) over 0–20 *µ*m, correlates with survival in PDAC (27), but has only been measured within individual 2D sections. Within the pathologist-annotated PDAC compartment, we computed the epithelial-to-T cell cross-L AUC for each 2D section and for the 3D volume, using H&E-predicted epithelial cells as the source population, and H&E-predicted T cells within the PDAC region and its surrounding 20 *µ*m margin as the target population (Fig. 5b–d, Methods). The 3D estimate exceeded every individual 2D section value: the mean section-wise AUC was 104.8*±*6.0 *µ*m^2^ across 40 serial sections, against 137.2 *µ*m^2^ for the volume, a 31% increase (Fig. 5g,h). This is because an epithelial-centered sphere captures T cells above and below the section plane that the corresponding 2D disc misses, and the discrepancy grows with the sphere radius (Fig. 5e, f). Since higher cross-L values reflect greater T cell proximity to epithelium, section-based estimates systematically overstate immune exclusion, raising the question of whether prognostic thresholds calibrated in 2D hold when measurements are made in the realistic 3D setting. We note that immune exclusion is quantitatively defined by the estimates being below the complete spatial randomness expectation (Fig. 5g).

We next examined whether the reconstructed cell maps could recover known differences in T cell–epithelial organization between high-grade IPMN and IPMN-associated PDAC. Multiplexed imaging of individual 2D sections has reported greater spatial association of CD8^+^ T cells with epithelium in IPMN than in IPMN-associated PDAC (40). Using pathologist-defined annotations, we delineated the two lesion compartments within the reconstructed pancreatic tissue and analyzed cells within each lesion and its surrounding 100 µm margin (Fig. 5i-k). To quantify local cellular organization in both 2D and 3D, we constructed Delaunay graphs (51) either independently within each section or across the SpaReg reconstructed volume. For each T cell, we then computed a class-specific interaction score as the fraction of its neighboring cells assigned to each cell class (Fig. 5l, Methods). When recomputed independently within each section, the mean T →Epithelial score was higher in IPMN than in PDAC throughout the annotated tissue depth (Fig. 5n). In both compartments, the observed scores were also below section-specific random-mixing expectations generated by label permutation, indicating T cell-epithelial exclusion in both lesions but to a greater degree in PDAC.

The volumetric analysis showed the same separation more strongly. The 3D T→Epithelial score maps showed greater epithelial association in IPMN than PDAC (Fig. 5m), with mean per-T-cell scores of 0.241 versus 0.065 (Fig. 5o). Conversely, T→other was higher in PDAC than IPMN (0.497 versus 0.334). Our classifier does not resolve the composition of this ‘other’ class, but in PDAC it is dominated by the desmoplastic stroma, comprising cancer-associated fibroblasts, activated stellate cells, dense extracellular matrix, and myeloid populations. The elevated T →other score in PDAC is therefore consistent with T cells residing within stroma rather than in contact with neoplastic epithelium, the pattern described for pancreatic stellate-cell sequestration of CD8^+^ T cells in the juxtatumoral compartment (28).

### 3D reconstruction resolves lymphoid aggregate continuity and tumor proximity

Beyond cell–cell interactions, 3D reconstruction also reveals the full spatial extent of multi-cellular immune structures. Organized lymphoid aggregates (LAs), particularly tertiary lymphoid structures (TLS), their more mature forms, are increasingly recognized as components of anti-tumor immunity. Across several cancers, TLS are associated with improved prognosis and immunotherapy response, reflecting their role as sites of organized T and B cell responses (24, 52), and in PDAC specifically, intratumoral and mature lymphoid structures have been linked to improved survival and tumor-specific immunity (30, 31). Both maturation state and position relative to tumor are prognostically informative (29), yet these features are typically assessed from individual sections, where apparent size, composition, and tumor proximity depend on the sectioning plane and 3D continuity is lost entirely. We therefore used cell-type-resolved SpaReg reconstruction to detect lymphoid aggregates across serial H&E sections and characterize their 3D spatial organization. Because canonical TLS classification requires markers beyond H&E, including follicular dendritic cell networks and high endothelial venules, we refer throughout to these structures as lymphoid aggregates rather than TLS.

Linking H&E-predicted T- and B-cell clusters across serial sections yielded ten 3D lymphoid aggregates, each spanning at least three sections (Methods). Supplementary Video 4 shows the reconstructed H&E tissue block from Fig. 6a, with its corresponding cell-type map, revealing the spatial organization of the PDAC and surrounding LAs in 3D. Across the 200 *µ*m thick reconstructed volume, individual aggregates changed substantially with depth, with structures prominent in one section appearing reduced, fragmented, or undetected in another (Fig. 6a). The aggregates ranged from 2,461 to 375,366 lymphocytes and differed in T and B cell composition, with B cells accounting for 1.0% to 22.15% of aggregate lymphocytes (Fig. 6b). This range is informative because B cell abundance is concordant with LA maturity, with early aggregates being predominantly T cell infiltrates, whereas mature TLS acquiring a B cell follicle surrounded by a T cell zone and are correspondingly B cell rich (24). We note that given our classifier’s B cell recall, the above proportions likely represent lower bounds on B cell content.

**Fig. 6.**
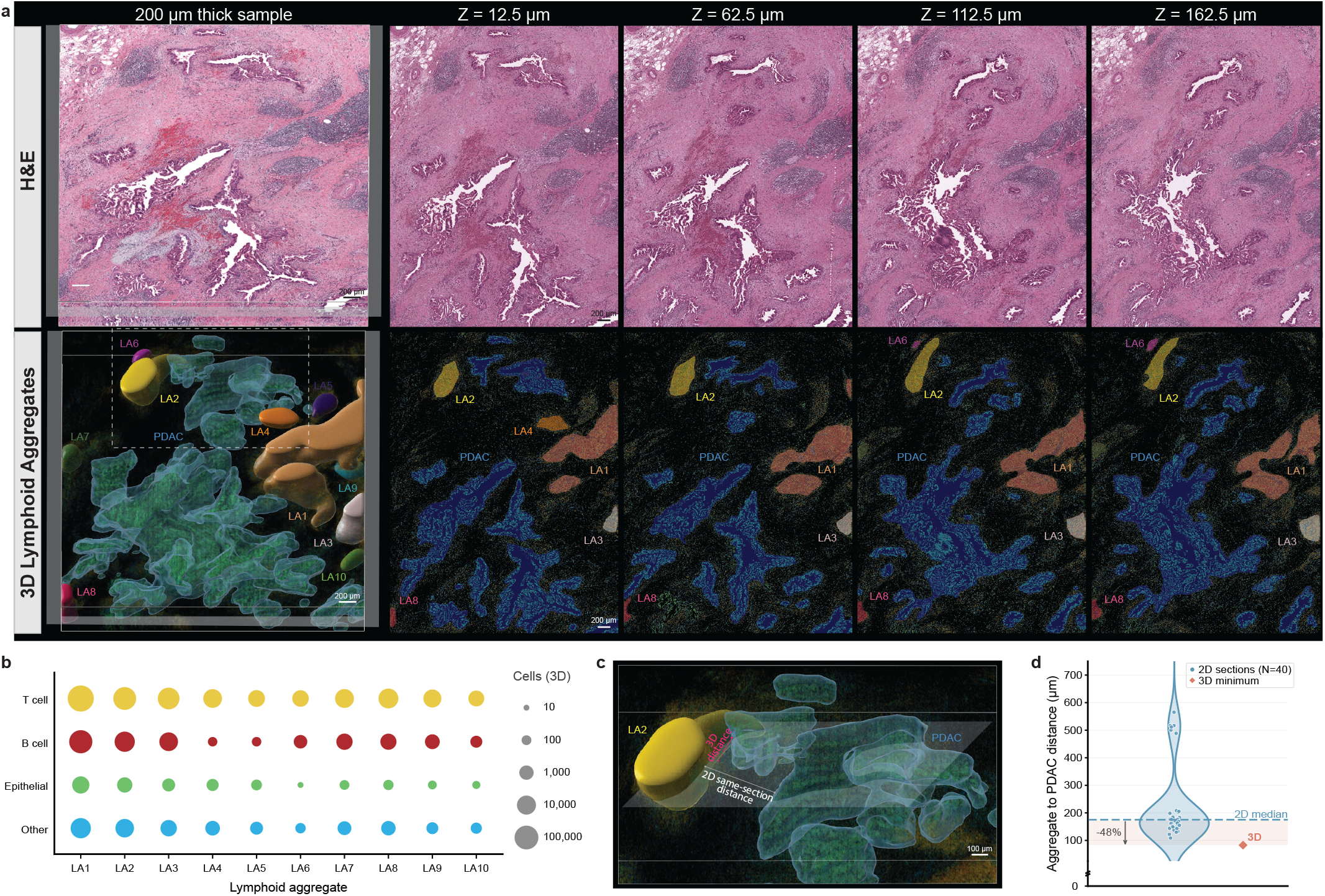
SpaReg resolves 3D lymphoid aggregates directly from H&E. **a** 3D H&E reconstruction of a 3.5 mm^3^ PDAC-containing tissue specimen and few 2D sections across tissue depth (top). Cell type map with resolved lymphoid aggregates LA1-LA10 and PDAC (bottom). See Supplementary Video 4. **b** Bubble plot of cell type composition of each 3D lymphoid aggregate. Bubble area is log-scaled to the number of cells assigned to each aggregate. **c** Illustration of aggregate proximity to PDAC, section-based 2D vs 3D. White line indicates the minimum same-section distance between LA2-associated lymphocytes and PDAC-associated epithelial cells; magenta line indicates the minimum 3D distance between the same populations. **d** Proximity of LA2-associated lymphocytes and PDAC-associated epithelial cells.

We then examined how 3D reconstruction changes the estimated proximity of lymphoid aggregates to PDAC. As an example, for aggregate LA2 (Fig. 6c), the median per-section minimum distance to PDAC epithelial cells was 168.5 *µ*m, against a 3D minimum distance of 87.1 *µ*m, a 48% reduction (Fig. 6d). This mirrors the geometric bias observed for epithelial–T cell proximity, showing that section-based measurements can underestimate spatial associations that occur across tissue depth.

## Discussion

SpaReg reframes serial-section reconstruction as alignment of sparse tissue-boundary coordinates, retrieving matched regions at the native image resolution. This geometric foundation supports H&E, multiplexed immunofluorescence and spatial transcriptomics, with subsequent refinement adapted to each modality. An immediate application is the construction of multi-modal 3D atlases linking gene expression, protein expression, and morphology across organs and disease states, so that molecular measurements can be interpreted in their architectural context and architectural features traced to their molecular basis.

A complementary opportunity lies in extending 3D microenvironment analysis to routine and archived histology. Our cell type maps, in which epithelial, T, and B identities are inferred throughout the reconstructed PDAC microenvironment from H&E alone, demonstrate this potential. As SpaReg-based 3D reconstruction works with standard WSIs from archived FFPE blocks, it can bring 3D spatial analysis into computational pathology, allowing spatial biomarkers such as intratumoral T cell density, immune exclusion, and lymphoid aggregate maturity, which are currently scored from single sections, to be scored in 3D. Our finding that 2D sections overstate immune exclusion suggests that prognostic thresholds calibrated in 2D may need re-evaluation in 3D before clinical use.

Beyond cell-type mapping, retaining cellular and subcellular morphology at the original image resolution provides a foundation for extending H&E-based prediction of spatial gene expression (53, 54), protein expression distributions (55, 56), and cell states (57) into 3D. Integrating these methods with SpaReg could enable molecularly annotated reconstructions for studying tumor heterogeneity and the spatial organization of immune and stromal niches. A hybrid design could support this extension by combining multiplexed imaging on a sparse subset of sections with H&E on the remainder, using measured signals to calibrate and evaluate molecular predictions. Such inference should complement direct molecular measurement and could reduce the reagent and instrumentation burden of characterizing 3D tissue microenvironments across larger series and cohorts.

Our results also show how spatial sampling affects cell-resolved measurements. Increasing the effective pixel size of classified cell maps merges adjacent cells, disproportionately removes lymphocytes, and shifts apparent composition. Dimensionality introduces a complementary sampling effect: the 3D tissue reconstruction captures relationships across tissue depth that individual sections miss, changing estimates of immune exclusion and aggregate proximity. These are systematic consequences of spatial sampling rather than random measurement noise and should therefore be considered when interpreting 2D and 3D spatial analyses. Integrating molecular measurements with reconstructed morphology could further allow the interacting cells and structures identified in 3D to be characterized by their molecular states.

Our classifier distinguished epithelial, T, and B cell identities, although lymphocyte classification was less accurate than epithelial classification in the eight-region training data. The compartment- and aggregate-level measurements reported above pool over many cells, but measurements that depend directly on the relative abundance of T and B cells remain sensitive to classification uncertainty. Extending training across additional markers and samples could improve classification performance and resolve additional cell classes. The paired H&E and IF imaging design used here can be extended to this setting without changing the overall framework.

## Methods

### Overview

SpaReg reconstructs tissue microenvironments in 3D from ordered serial whole-slide images or spatial transcriptomics spot or cell coordinates. Image inputs include hematoxylin and eosin (H&E), multiplexed immunofluorescence (IF), and cross-modal H&E–IF serial sections.

SpaReg follows two design principles. First, initial registration uses sparse tissue-boundary geometry rather than dense image or molecular content, providing a common geometric basis for alignment across modalities. Second, image-based reconstruction propagates region-of-interest (ROI) coordinates rather than warping entire whole-slide images. Regions corresponding to the ROI across the serial sections are extracted from their respective WSIs at the native image resolution. Any residual misalignment between these regions is corrected via recursively improving coordinate-mapping.

For image-based 3D reconstruction, the user defines the ROI once on any selected serial section, which serves as the anchor and establishes the common coordinate frame. SpaReg automatically registers the ROI outward from this anchor section across the set of ordered serial sections. For spatial transcriptomics, the estimated transformations are applied directly to spot or cell coordinates, with optional point-cloud refinement to accommodate residual geometric differences.

The modality-specific implementations and parameter choices are described below (Fig. 1b and Extended Data Fig. 1). Datasets and software versions are listed in Supplementary Tables 1 and 2, respectively. Registration parameters were held fixed across all reconstructions and benchmark studies included in this manuscript, unless otherwise explicitly specified for a given modality.

### SpaReg 3D reconstruction of image-based tissue sections

#### Automated ordering of serial sections from slide labels

WSI filenames of individual serial sections do not always preserve the physical order of serial sections, particularly when slides are rescanned or renamed during institutional or laboratory quality-control workflows. The original section number is generally retained in the slide-label image embedded within the WSI. When needed, SpaReg uses this information to recover the physical section order independently of the file-name.

For each WSI, the associated label image is extracted using OpenSlide (58), and optical character recognition is performed using EasyOCR (59) with the English-language recognition model. Labels are converted to grayscale, binarized using Otsu (60) thresholding, and re-analyzed to recover faint or low-contrast digits.

The section-number field is identified jointly across all serial sections rather than from a predefined label template. Numerical detections are grouped by their spatial position on the label. Fields that remain constant or repeat across slides, including patient identifiers, anatomic site, institution or laboratory name, and accession identifiers, are treated as fixed label content and excluded. The remaining numerical field that appears at a consistent label position but varies across slides is retained as the section number. For each slide, the highest-confidence detection from this field is parsed as the section index, and the sections are ordered accordingly for 3D reconstruction.

#### Tissue-foreground extraction

For image-based reconstruction, a binary tissue foreground is obtained from each section. Its external contour defines the tissue boundary used for coarse registration. The foreground-extraction strategy depends on the imaging modality.

##### Hematoxylin and eosin

For H&E sections, SpaReg segments tissue using local chroma entropy rather than grayscale intensity, allowing heterogeneous and weakly stained tissue to be retained in the foreground mask (Extended Data Fig. 1b–d, Extended Data Fig. 8). Each downsampled whole-slide image is converted to the CIELAB color space, and a chroma image is derived from the two chromatic channels and normalized to an 8-bit range. The local Shannon entropy of the chroma image is computed within a small disk-shaped neighborhood around each pixel. The resulting entropy map is thresholded using the more conservative of the mean entropy and the Otsu-derived threshold to obtain the tissue mask. Because chroma entropy reflects local variation in tissue color and texture rather than absolute intensity, weakly stained compartments such as fibroadipose tissue are retained more reliably than with intensity-based thresholding alone.

##### Multiplex immunofluorescence

For IF sections, the tissue foreground is derived from the Hoechst/DAPI channel, which provides a common structural signal across the serial sections. The Hoechst/DAPI image from each section is down-sampled and binarized using Otsu thresholding.

##### Cross-modal H&E and IF

For cross-modal reconstruction of interleaved H&E and IF sections from the same specimen, tissue foregrounds are generated using the modality-specific procedures described above. H&E sections therefore use chroma-entropy-based foreground extraction, whereas IF sections use the Hoechst/DAPI signal. For subsequent cross-modal refinement, the hematoxylin component of H&E and the Hoechst/DAPI channel of IF provide a shared nuclear signal, as described below.

#### Sparse tissue-boundary representation

The external tissue contour is extracted from the binary foreground mask (Extended Data Fig. 1e) and lightly smoothed to suppress small boundary irregularities. An *α*-shape is then constructed from the contour points, preserving concavities in the tissue outline that would be lost with a convex-hull representation. The *α* parameter is 0.001 for H&E and 0.03 for IF, and the contour-smoothing parameter *σ* is 1.

The resulting boundary is reduced to a sparse set of representative keypoints by *k*-means++ clustering (61). The boundary representation uses at most 1000 keypoints. Each cluster center is then mapped to its nearest observed contour point so that every retained keypoint remains on the tissue boundary. This sparse representation retains the overall tissue geometry while reducing the number of points used for subsequent boundary matching. Since the sparse correspondence is distributed across the tissue boundary, localized sectioning artifacts that disrupt internal tissue or portions of the tissue boundary leave sufficient intact geometry for SpaReg alignment to be robust to them (Extended Data Fig. 1a, tear, Extended Data Fig. 2a, fold).

We quantify sparsity as *s* = *K/n*, where *K* is the number of keypoints and *n* is the number of tissue pixels in the section at native resolution. We count *n* at native resolution because the estimated transformation is applied in native coordinates and the reconstruction is delivered at that resolution, whereas downsampling serves only to extract the tissue contour. For the H&E series in this study, *K* = 1000 and *n* = 1.8–8.5 *×* 10^9^ (from the section sizes and resolutions in Supplementary Table 1), giving *s≈* 1–4 *×* 10^*−*7^, that is, 0.00001–0.00004% of the pixels in a section.

#### Rotation-invariant boundary matching

Boundary keypoints from the *α*-shape representation are described using rotation-invariant shape-context descriptors (Extended Data Fig. 1f,g). For each keypoint, the descriptor summarizes the angular and radial distribution of all other boundary key-points, with angles measured relative to the direction from the tissue centroid to the keypoint. This allows corresponding boundary geometry to be compared independently of the global section orientation. Descriptor dissimilarity is quantified by the chi-square distance, and one-to-one correspondences are obtained by solving the minimum-cost assignment between the two keypoint sets using the Hungarian algorithm (62, 63) (Extended Data Fig. 1h).

High-cost correspondences are removed as outliers using Tukey filtering before coarse transformation estimation. For cross-modal pairs, the coarse orientation is additionally resolved using a mutual-information search over rotation. This provides a suitable similarity measure when the corresponding H&E and IF sections have different intensity distributions.

#### Coarse contour registration

The retained correspondences are used to estimate a two-dimensional rigid transformation between the tissue boundary of the section being registered (moving section) and that of the reference section. Rotation and translation are obtained in closed form by solving the orthogonal Procrustes problem with the Kabsch solution (64), constrained to proper rotations so that reflected solutions are excluded. The resulting transformation maps the moving-section coordinates into the reference coordinate frame.

The transformation is then refined iteratively. Correspondences with large spatial residuals are progressively removed, and the rigid transformation is re-estimated from the remaining matches. Refinement is retained only when tissue-mask overlap is maintained or improved. When residual misalignment remains, iterative closest-point matching (ICP) (65) is applied to the complete tissue boundaries. At this stage, the sections are already approximately aligned, allowing correspondence to be refined by spatial proximity. Distant point pairs are removed as outliers before re-estimating the rigid transformation. If a moving section cannot be aligned reliably, it is not used as the reference for the next pairwise registration. Registration instead continues from the most recent well-aligned reference section, limiting propagation of alignment error through the serial sections. For IF, reference propagation requires a contour Dice score of at least 0.30.

As this stage estimates rotation and translation, it aligns global tissue geometry without introducing scaling, shear, or local deformation. Tissue regions present in one section but absent from an adjacent section therefore retain their original shape rather than being stretched or compressed to create a correspondence (Extended Data Fig. 1i). This preserves section-specific morphology before subsequent full-resolution refinement.

#### ROI coordinate mapping and image retrieval at native resolution

Following coarse contour registration, the user-defined region of interest (ROI) is mapped from the reference section to its corresponding location in each moving section. Rather than transforming the entire whole-slide image, SpaReg maps the vertices of the ROI polygon into the moving-section coordinate frame using the inverse of the coarse transformation (Extended Data Fig. 1j).

The transformed polygon coordinates are converted from the downsampled registration space to the native image resolution. The corresponding image region enclosing the mapped polygon is then retrieved directly from the original whole-slide image of the moving section (Extended Data Fig. 1k). This preserves the subcellular morphological detail present in the acquired image.

For a whole-slide image of dimensions *N×N*, conventional whole-image warping requires resampling *N*^2^ pixels and therefore scales as *O*(*N*^2^). SpaReg instead transforms only the *p* vertices defining the ROI polygon, giving *O*(*p*) coordinate-mapping complexity. Because *p* is fixed with respect to whole-slide image dimensions, this operation is *O*(1) with respect to *N*. The subsequent image read scales with the *A* pixels enclosing the mapped ROI rather than with the full *N*^2^ image, where *A≪ N*^2^ for a localized tissue region.

This separation of coordinate mapping from image retrieval allows regions within gigapixel whole-slide images to be accessed at their native image resolution without whole-slide resampling. Because the mapped region is retrieved directly from the original image, its original pixel values and subcellular morphological detail are retained for subsequent refinement.

#### Refinement at native image resolution

Coarse contour registration localizes corresponding tissue regions using boundary geometry at downsampled resolution. After retrieval at the native image resolution, the internal tissue signal, including cellular and subcellular morphology, provides an additional source of correspondence for local refinement (Extended Data Fig. 1l).

The moving and reference ROIs are aligned by maximizing the enhanced correlation coefficient (ECC) (66) over the shared tissue foreground, estimating a rigid transformation first and then refining it to an affine transformation. This corrects the residual translation, rotation, and small differences in local scale that remain after contour registration. A refinement is retained only when it improves the intensity correlation between the ROIs.

For same-modal reconstruction of serial H&E or serial IF sections, ECC operates directly on the ROI intensities. For cross-modal pairs, refinement uses the shared nuclear signal, with the hematoxylin component for H&E and DAPI for IF. After refinement, the updated ROI coordinates are applied to the original whole-slide image and the corresponding region is retrieved again at the native image resolution. This retains the original pixel values and subcellular morphological detail in the final aligned region.

#### Elastic refinement

Elastic refinement is an optional final step. Across the serial H&E, immunofluorescence, and cross-modal reconstructions in this study, spanning multiple tissue types and hundreds of serial sections, the native-resolution refinement already corrects the residual misalignment left by contour registration and brings adjacent sections into near-perfect alignment. The additional elastic step, which is considerably more computationally expensive, is therefore seldom necessary.

When applied, residual local deformation between adjacent sections is corrected by bounded symmetric diffeomorphic registration (SyN) (67). The reference and aligned moving ROIs are converted to grayscale before SyN registration. The displacement cap is 60 pixels, corresponding to 15 *µ*m at 0.25 *µ*m/px. The estimated displacement field is accepted only when it increases the normalized cross-correlation between the reference and moving ROIs. Otherwise, the transformation from the preceding rigid or affine refinement is retained.

### SpaReg 3D reconstruction of spatial transcriptomics data

SpaReg registers spatial transcriptomics (ST) sections using the same contour-based geometric framework as image-based serial sections. The tissue outline is constructed directly from measured spot or cell coordinates rather than from an image-derived foreground. Sparse boundary representation, rotation-invariant shape-context matching, Hungarian assignment, and coarse transformation estimation are applied directly to these coordinate sets as described above for image-based reconstruction. Registration therefore operates on the measured spatial point sets without converting gene-expression measurements into images. Where residual geometric differences remain after coarse alignment, SpaReg applies an optional deformable point-cloud refinement.

#### Tissue-outline extraction and boundary matching

The tissue outline is constructed directly from the measured spot or cell coordinates. For regularly sampled Visium spot grids, a convex hull captures the overall tissue extent. For single-cell point sets such as MERFISH, an *α*-shape preserves concave anatomical features of the tissue boundary, including fissures and ventricular boundaries. The exterior boundary is resampled at uniform arc length and lightly smoothed before correspondence matching, with contour-smoothing parameter *σ* = 3.

Boundary correspondences are obtained using the geometric matching framework described above for image-based reconstruction, with rotation-invariant shape-context descriptors, chi-square descriptor distance, and one-to-one Hungarian assignment.

Because ST sections can differ substantially in orientation, candidate correspondences are filtered using a RANSAC-style geometric-consistency search (68). Candidate transformations are estimated from subsets of correspondences, and the transformation supported by the largest geometrically consistent set is retained. The inlier radius is 0.05 times the bounding-box diagonal. This allows valid boundary correspondences to be identified across large differences in section orientation.

#### Transformation estimation

Rigid, similarity, and affine candidate transformations are estimated from the retained boundary correspondences for each section pair. Similarity transformations are restricted to near-isotropic scaling, while affine transformations are constrained to limit anisotropic deformation. Candidate rotations are evaluated over the full 0–360^*°*^ range about the tissue centroid, followed by a finer angular search around the orientation with the best boundary agreement. Reflected candidates are evaluated in parallel when section mounting could introduce a left–right reversal. Candidate rotations, reflections, and transformation models are ranked using symmetric Chamfer distance, defined as the mean nearest-neighbor distance between the reference and transformed moving boundaries in both directions, with lower values indicating closer geometric agreement. For geometrically ambiguous sections, particularly bilaterally similar anatomy, candidate selection additionally uses cell-type-restricted Chamfer distance, computed separately for each represented cell-type class and averaged across classes. This favors alignment of corresponding cellular compartments when the outer boundaries alone cannot distinguish candidates. Cell-type labels are used only for candidate selection and do not define point-wise correspondences or serve as registration targets. Anatomical region labels used for benchmarking were not used during registration.

The selected transformation is refined using the most spatially consistent correspondences and rigid iterative closest-point (ICP) matching of the complete tissue boundaries. During ICP refinement, distant point pairs are removed before re-estimating the rigid transformation, and the refinement is retained only when it improves the symmetric Chamfer score.

#### Non-rigid point-cloud refinement

Non-rigid point-cloud refinement is an optional final step applied when residual local geometric differences remain after rigid or affine alignment. In this study, it is used for geometrically complex reconstructions such as the 129-section MERFISH dataset.

Residual boundary differences are modeled using deformable Coherent Point Drift (CPD) (69), which models the moving points as a Gaussian mixture and estimates coherent, neighborhood-preserving displacements to the reference. CPD operates on sampled boundary points after the moving and reference point sets are placed in a common normalized coordinate space. The resulting boundary displacements are propagated through the tissue interior using thin-plate-spline interpolation. This converts the boundary displacement estimated by CPD into a smooth deformation field that is applied to the complete spatial point set. Molecular measurements and annotations remain associated with their corresponding transformed spots or cells.

### H&E cell type classifier development and training

#### Cyclic immunofluorescence staining and imaging

For training the cell classification model, we performed cyclic immunofluorescence (CyCIF) (9, 42) imaging on a pancreatic tissue section, containing PDAC, to derive cell-type ground truth from protein expression. Specifically, a deidentified formalin-fixed paraffin-embedded (FFPE) tissue section containing PDAC was obtained from the Department of Pathology at the University of Pittsburgh, and processed using the CyCIF antigen retrieval protocol (9, 42). The slide was stained over CyCIF cycles, with three antibodies and DAPI acquired per cycle. The cell type classifier panel used E-cadherin (CST 3199S, clone `24E10`, 1 in 100), pan-cytokeratin (Invitrogen 41-9003-82, clone `AE1/AE3`, 1 in 50), CD4 (eBioscience 41-2444-82, clone `N1UG0`, 1 in, CD8a (Invitrogen 50-0008-82, clone `AMC908`, 1 in 20), CD20 (CST 83399S, clone `E7B7T`, 1 in 100), and CD79a (Novus NB100-64347AF488, clone `HM57`, 1 in 100). Antibody staining was performed overnight at 4^*°*^C in the dark. Each cycle was followed by staining with Hoechst 33342 (CST 4082S) for 10 min at room temperature in the dark.

Images were acquired after each cycle with a 20X*/*0.75NA objective on a Nikon Ti2E microscope at 0.32 *µ*m/pixel. Eight high-quality regions were identified across the slide and extracted for classifier training.

#### H&E staining and cell segmentation

Following CyCIF imaging, the same pancreatic tissue section was H&E stained and imaged on a Leica Aperio AT2 whole-slide scanner at 40X magnification and 0.25 *µ*m/px (Supplementary Table 1). H&E regions corresponding to the eight CyCIF acquisitions were identified and extracted from the whole-slide image. Nuclei were detected and segmented using CellViT++ (39) with the Virchow histopathology foundation model (43) as its encoder, selected for its zero-shot segmentation performance across diverse histopathology datasets. We used only the segmentation output, bypassing the built-in classification head, to obtain region-wise nuclear boundaries, which were converted into nuclear instance masks. This yielded 181,976 segmented nuclei across the eight regions (Fig. 1c–e).

To associate the H&E nuclear instances with their corresponding CyCIF marker measurements, each H&E region was registered to its matched CyCIF image. The H&E grayscale image was resampled to the CyCIF resolution of 0.32 *µ*m/px by bicubic interpolation and the instance mask by nearest-neighbor interpolation, preserving discrete cell identities. The estimated transform was applied to the instance mask, producing a registered H&E nuclear-instance mask in the CyCIF coordinate frame.

#### CyCIF cell profiling and cell-type assignment

Images obtained from successive CyCIF staining cycles were coregistered using Hoechst nuclear signal as the common reference. Registration was performed using SpaReg, and the resulting transformations were applied to each channel of each CyCIF cycle. Non-Hoechst marker images were also corrected for spatial variation in illumination using the BaSiC flat-field correction algorithm (70), followed by rolling-ball background subtraction (71, 72).

To associate CyCIF marker measurements with the corresponding H&E cell instances, whole-cell segmentation of the CyCIF images was initialized from the registered H&E-derived nuclear instance masks using UNSEG (73). Each resulting whole-cell segmentation therefore retained the identifier of its corresponding H&E nuclear instance. Marker intensities were normalized by illumination power and exposure time, and cell-level expression is defined as the mean normalized intensity across the pixels of each whole-cell segmentation.

Cell types were assigned from the CyCIF-derived marker profiles using a rule-based procedure, with pan-cytokeratin and E-cadherin defining epithelial cells, CD4 and CD8a defining T cells, and CD20 and CD79a defining B cells. For each marker, cell-level expression was winsorized and min-max normalized. Marker positivity was then defined using marker-specific thresholds (Supplementary Table 3). The median was used for pan-cytokeratin and E-cadherin, triangle thresholding (74) for CD4, CD8a and CD20, and the 0.98 empirical quantile for CD79a. Sub-threshold values were set to zero, and a score was computed for each class by summing the retained intensities of its defining markers. Cells were assigned to the highest-scoring class, with near-ties resolved by the single most strongly expressed marker. Cells without sufficient marker support for any class were assigned to the ‘other’ category.

Image quality control identified two regions with elevated nonspecific signal in the CD20 channel. B cell assignment in these regions used more stringent thresholds: a log-Otsu threshold for CD20 and the 0.995 empirical quantile for CD79a (Supplementary Table 3). B cell assignments judged inconsistent with the corresponding CyCIF images during image review were reassigned to the next highest-scoring class.

#### H&E patch generation and classifier training

We mitigated the effect of H&E stain variation due to CyCIF by applying Vahadane stain normalization (75) to the eight annotated PDAC regions, using an H&E-stained whole-slide image of pancreatic tissue that had not undergone CyCIF imaging, as the reference. Each normalized region was divided into 256 *×* 256 pixel image patches with 10% overlap between adjacent patches, yielding 12,962 patches and 224,657 cell-label instances from the 181,976 unique segmented nuclei. To increase representation of the less abundant lymphocyte classes during training, we sampled additional patches around T- and B-cell centroids with randomized spatial offsets, while limiting excessive overlap between patches.

For evaluating the cell classification model, we used four-fold cross-validation at the region level so that spatially overlapping patches from the same tissue region could not appear in both training and validation sets. The eight PDAC regions were partitioned into four non-overlapping pairs, so that within each fold, six regions could be used for training, and the remaining two held out for validation, with each region appearing in validation exactly once (Fig. 4a,b). Training sets included both the regularly sampled and additional lymphocyte-centered patches, whereas validation used only the regularly sampled patches from held-out regions.

The cell classification model was trained within the CellViT++ framework (39) using the Virchow histopathology foundation model (43) as a frozen encoder and classifies each detected cell as epithelial, T cell, B cell or Other. Virchow-derived embeddings associated with each H&E cell instance were passed to a multilayer perceptron classifier with Gaussian Error Linear Unit (GELU) activations (76) and a dropout probability of 0.5. Model parameters were optimized using AdamW (77) with *β*_1_ = 0.85, *β*_2_ = 0.9, a learning rate of 2 *×*10^*−*5^, and weight decay of 0.08. To account for differences in class abundance, the cross-entropy loss was weighted by the inverse square root of the frequency of each class. Models were trained with a batch size of 128 for up to 50 epochs, with early stopping after 10 consecutive epochs without improvement in the validation macro F1 score.

Model performance was summarized across the four heldout folds, with per-class and macro F1 scores calculated independently within each fold from the predicted class with maximum probability and reported as the mean *±* s.d. across folds. Confusion matrices were row-normalized within each fold before averaging, with each entry reported as the across-fold mean *±* s.d. Following cross-validation, a final classifier was trained using patches from all eight annotated PDAC regions for application to additional H&E sections, with 7% of the regularly sampled patches from each region held out for early stopping.

### SpaReg benchmarking and evaluation

#### Benchmarking 3D reconstruction accuracy and tissue integrity

To benchmark 3D reconstruction accuracy and tissue integrity, we used a publicly available dataset comprising 260 serial H&E-stained sections of murine prostate generated and published by Kartasalo et al. (16). The sections were acquired at 0.46 *µ*m/px, with a section thickness of 5 *µ*m.

The benchmark includes manually annotated corresponding cellular landmarks between adjacent sections from two independent researchers. Landmark displacement was used to calculate accumulated target registration error (ATRE), which quantifies registration error propagated through the serial sections and therefore provides a measure of 3D reconstruction accuracy.

Methods included in Fig. 3a are OPT, SIFT (78), HyperStackReg (HSR) (72), RegisterVirtualStackSlices (RVSS) (79), ElasticStackAlignment (ESA) (80), Medical Image Manager (MIM), Voloom, CODA (14), VALIS (15), and SpaReg. Where both default and optimized parameter settings are reported, the optimized configuration was used. The unregistered sections served as a no-registration reference, and least-squares registration using the published landmarks (LS1) served as a landmark-informed reference.

Published ATRE and tissue-area-change measurements were obtained from the original benchmark (16) and, for CODA, from its published evaluation (14). Because corresponding measurements are not reported for VALIS, VALIS and SpaReg were evaluated on the same serial sections using the original RegBenchmark implementation together with the published landmark annotations and tissue masks. SpaReg was evaluated after coarse contour registration, denoted SpaReg coarse, and after refinement at the native image resolution, denoted SpaReg fine.

3D reconstruction accuracy was quantified using ATRE, which measures the cumulative displacement of corresponding landmarks through the registered serial sections. ATRE therefore captures alignment errors that may be small between adjacent sections but accumulate through the reconstructed tissue. Lower ATRE indicates greater 3D reconstruction accuracy. For Fig. 3a, ATRE was converted to the higher-is-better *Accumulated Target Registration Accuracy* score shown on the x-axis.

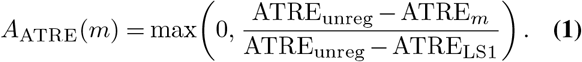

Here, *m* denotes the reconstruction method. Using the reference values reported in the original benchmark (16), ATRE_unreg_ = 1153.1 *µ*m and ATRE_LS1_ = 3.6 *µ*m. The resulting score represents improvement from the unregistered sections toward the landmark-informed reference, with values below zero set to zero. Higher Accumulated Target Registration Accuracy therefore indicates lower cumulative landmark displacement and greater 3D reconstruction accuracy.

Tissue integrity was quantified from the percentage change in tissue-mask area introduced by registration, *dA*. Positive values indicate tissue expansion, negative values indicate contraction, and values near zero indicate preservation of the original section area. Because both expansion and contraction represent geometric alteration, the magnitude of the mean area change was converted to the higher-is-better *Tissue Integrity Score* shown on the y-axis of Fig. 3a.

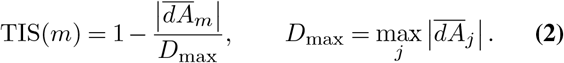

Here, 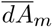 is the mean percentage tissue-area change for method *m*, and *D*_max_ is the largest absolute mean area change among the methods included in the benchmark. For the methods evaluated here, *D*_max_ = 33.10%. A Tissue Integrity Score of 1 indicates preservation of the original tissue area, whereas 0 represents the largest geometric alteration observed in this comparison. Accumulated Target Registration Accuracy and Tissue Integrity Score are plotted jointly in Fig. 3a to assess reconstruction accuracy together with preservation of tissue geometry.

#### Benchmarking spatial transcriptomics 3D reconstruction

Spatial transcriptomics 3D reconstruction was evaluated using the published SABench framework (22) on the datasets represented in Fig. 3c–d. These comprise four serial 10x Visium sections of human dorsolateral prefrontal cortex (DLPFC) and five MERFISH sections of the mouse hypothalamic preoptic region (MHPR). SpaReg was applied to the same spatially perturbed inputs used in SABench, and its outputs were evaluated using the same marker-gene sets, anatomical annotations, and metric definitions provided by the benchmark. Published measurements for the other methods were taken from the corresponding SABench results.

Methods shown in Fig. 3c–d include PASTE (10), PASTE2 (17), STAligner (21), GPSA (20), SLAT (18), STalign (11), CAST (81), STAIR (82), SPACEL (83), and Spateo (12), depending on availability for each dataset. Where SABench reports separate configurations that are also shown separately in the figure, these were retained as distinct benchmark entries, including rigid and non-rigid Spateo, shown as Spateo-r and Spateo-nr, and PASTE-p0.

The x-axis of Fig. 3c–d, *Gene expression similarity*, summarizes agreement of spatial gene-expression patterns between reconstructed sections. Following SABench, markergene expression was compared across corresponding spatial regions of aligned section pairs using the Pearson correlation coefficient (PCC), cosine similarity, structural similarity index (SSIM), and mutual information (MI). These four measurements were averaged to obtain the composite Gene expression similarity score.

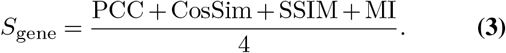

Higher values indicate greater concordance of spatial gene-expression patterns between reconstructed sections.

The y-axis, *Spatial landmark accuracy*, summarizes anatomical agreement between reconstructed sections. SABench evaluates this using matching accuracy of anatomical annotations together with spatial overlap of corresponding anatomical regions. These two measurements were averaged to obtain the composite Spatial landmark accuracy score.

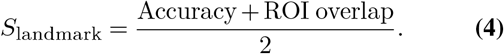

For DLPFC, the anatomical annotations correspond to cortical layers and white matter. For MHPR, they correspond to the annotated anatomical regions used in SABench. Higher Spatial landmark accuracy indicates greater anatomical correspondence between reconstructed sections.

#### Additional evaluation of spatial transcriptomics reconstruction

To quantify accumulation of spatial error through the DLPFC reconstruction, we measured the displacement of shared cortical-region centroids between consecutive registered sections. For each section pair, the mean displacement of anatomical regions present in both sections was calculated. To account for differences in coordinate scale, this displacement was normalized by a characteristic tissue radius, defined from the root-mean-square distance of spots from the centroid of each unregistered section and summarized by the median across sections.

Accumulated reconstruction error through section *k* was calculated as

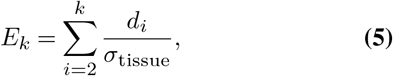

where *d*_*i*_ is the mean displacement of shared anatomical-region centroids between consecutive sections and *σ*_tissue_ is the median tissue radius. Accumulated error is reported in tissue-radius units, with lower values indicating less spatial drift through the reconstructed tissue.

To evaluate whether differences in reconstruction accuracy affect downstream spatial analysis, we adapted the DLPFC 3D spatial-domain clustering workflow from SABench (22). Four adjacent 10x Visium sections from donor III (151673– 151676) were used, with expert annotations for cortical layers 1–6 and white matter providing seven reference spatial domains. Each section was independently perturbed by rotation, translation, and reflection before reconstruction. SpaReg was evaluated alongside PASTE2 (17), SPACEL (83), CAST (81), rigid and non-rigid Spateo (12), and an unaligned baseline.

The same expression measurements and feature-processing workflow were used for every method, with only the reconstructed spatial coordinates differing between analyses. GraphST (84) was applied to the reconstructed sections to obtain a joint spatial representation. Following the SABench workflow, the GraphST embedding was reduced to 20 principal components and partitioned into seven spatial domains using `mclust` (85) with the EEE covariance model.

Predicted spatial domains were compared with the expert cortical-layer annotations separately within each section using the adjusted Rand index (ARI), normalized mutual information (NMI), homogeneity (HOM), and completeness (COM). The four measurements were averaged across the four sections to obtain the overall downstream spatial-domain score.

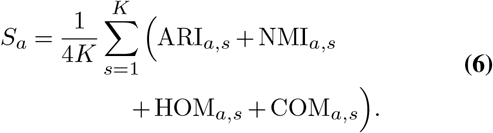

Here, *K* = 4 is the number of DLPFC sections and *s* indexes the individual sections. Higher values indicate closer agreement between the inferred spatial domains and the expert cortical-layer annotations.

#### Large-scale spatial transcriptomics 3D reconstruction benchmark

Large-scale spatial transcriptomics reconstruction was evaluated using the Zhuang-ABCA-1 whole-mouse-brain MERFISH dataset comprising 129 serial sections and approximately 2.64 million cells (Supplementary Table 1). To create a reconstruction challenge with known orientation ground truth, each section was independently rotated in plane by a known angle *θ*_*i*_ before reconstruction. No scaling was introduced.

SpaReg was evaluated alongside SPACEL (83), PASTE2 (17), CAST (81), and rigid and non-rigid Spateo (12). These methods were selected from the strongest-performing methods in the published SABench evaluations relevant to this study (22). PASTE2 and SPACEL rank among the leading methods on sequencing-based datasets, Spateo achieves the highest overall scores on imaging-based datasets, and CAST is among the top-performing methods in downstream 3D spatial-domain analysis. The independently rotated input was retained as the unaligned reference.

Each method was applied to the complete set of 129 sections using all measured cells and the method-specific workflow described in its corresponding publication. Successful completion was defined as returning reconstructed spatial coordinates for all 129 sections. Methods that terminated because of an execution error, insufficient memory, or failure to return a complete reconstruction were recorded as not completed.

Orientation recovery was quantified by comparing the known rotation applied to each section with the rotational component recovered during reconstruction. For each section, the rotational transformation between the perturbed input coordinates and the reconstructed coordinates was estimated using a two-dimensional Kabsch fit after centering both coordinate sets. Because the orientation of the 3D tissue reconstruction is defined only up to a common global rotation, the shared orientation across sections was estimated from the circular mean of *ϕ*_*i*_ + *θ*_*i*_ and removed before calculating section-specific residual error.

For section *i*, the residual orientation error is defined as

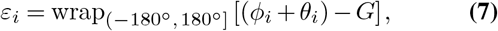

where *θ*_*i*_ is the known rotation applied before reconstruction, *ϕ*_*i*_ is the rotation recovered from the reconstructed coordinates, and *G* is the common global orientation estimated across the 129 sections.

Orientation recovery was summarized by the median absolute rotation residual and the fraction of sections with |*ε*_*i*_| *>* 15^*°*^, corresponding to the measurements shown in Extended Data Fig. 4. Lower values indicate more accurate and consistent recovery of section orientation across the reconstructed tissue in 3D.

### Spatial analyses of SpaReg-reconstructed tissue microenvironments

The analyses below were performed on 3D reconstructions of pancreatic tissue, using cell identities predicted from H&E by the classifier described above.

#### Three-dimensional cell detection

3D cell counts were obtained by aggregating 2D cell instance masks and their centroids within the 3D reconstructed volume using the spot-detection module in Imaris (11.0.0, Oxford Instruments), with an estimated spot diameter of 2.5 *µ*m and background subtraction disabled. These settings were held fixed across every analysis reporting Imaris-derived counts and across all compared conditions within each analysis, so that differences between resolutions, compartments, and dimensionalities reflect the underlying cell maps rather than the detection configuration.

#### Resolution degradation and cell retention

To quantify how image resolution affects the preservation of individual cell instances, we used the 320-section PDAC-containing reconstruction at the native H&E resolution of 0.25 *µ*m/px. Cells were segmented and classified on each registered section at native resolution, and the resulting boundaries and predicted cell types were rasterized into section-wise labeled instance masks. Resolution degradation was then simulated by down-sampling these masks to effective pixel sizes of 0.5, 1.0, 2.0, and 4.0 *µ*m/px by nearest-neighbor interpolation. Segmentation and classification were not repeated at the degraded resolutions, isolating the effect of spatial resolution on cell-instance separability from any change in classifier behavior.

Cell centroids detected at native resolution provided the reference, one entry per detected cell. At each degraded resolution, spatially separable cells were identified by connected-component analysis with 4-connectivity. Where several original instances merged on downsampling, the resulting component was counted as a single retained object and assigned a cell type by majority vote among the labels of its contributing pixels.

Within individual sections, retained cell counts were calculated independently for each serial section, cell type, and effective resolution, and cell retention was defined as the number of retained objects at a given resolution divided by the corresponding count at 0.25 *µ*m/px. Retention across the 320 sections was summarized by its median and interquartile range (IQR) for each cell type and resolution.

For 3D analysis, retained cell maps at each resolution were assembled into cell-type-specific volumes and quantified by spot detection in Imaris using the settings described above. 3D cell retention was defined analogously, as the cell count across the reconstructed tissue at a given downsampled resolution relative to the native H&E resolution of 0.25 *µ*m/px. To establish whether resolution loss also alters the apparent cellular composition of the tissue, we calculated the fraction of retained cells belonging to each cell type at each resolution and reported its change, in percentage points, relative to the native H&E resolution of 0.25 *µ*m/px (Fig. 4c,d).

#### Two- and three-dimensional L-function analysis

To quantify spatial association between epithelial cells and T cells in PDAC, the two populations were represented as the components of a bivariate spatial point pattern and their relationship characterized by the bivariate L-function, a transformation of the bivariate extension of Ripley’s K-function (49, 50). Following Carstens et al. (27), we summarized this association by the area under the L-function curve (AUC) over 0– 20 *µ*m, and extended the analysis by comparing estimates from individual 2D sections with the corresponding SpaReg-reconstructed tissue in 3D.

The analysis used a PDAC-containing ROI spanning 40 serial sections. The PDAC compartment within the ROI was manually annotated in each section and reviewed and confirmed by a pathologist. H&E-predicted epithelial cells with centroids inside the pathologist-confirmed PDAC region formed the source population, and H&E-predicted T cells within that region and a surrounding 20 *µ*m margin the target population. Margins were generated independently in each section by morphological dilation of the binary PDAC annotation by 20 *µ*m in the section plane.

For the 2D analysis, the epithelial-to-T-cell cross-L function was calculated independently within each serial section. For section *s*, let *E*_*s*_ denote the epithelial source coordinates, *T*_*s*_ the T cell target coordinates, and *A*_*s*_ the area of the PDAC analysis window including its margin. For each radius *r*,

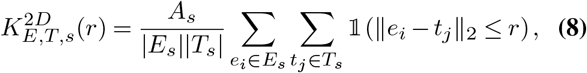

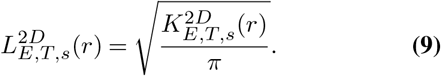

The cross-L function was evaluated at 1 *µ*m intervals from 1 to 20 *µ*m, with *L*(0) = 0 included explicitly, and its AUC over 0–20 *µ*m obtained by trapezoidal integration.

For the 3D analysis, epithelial and T cell maps from the 40 registered sections were assembled into 3D volumes and cell coordinates in three dimensions were obtained by spot detection in Imaris using the settings described above. The analysis volume was estimated from the section-wise PDAC windows using the Cavalieri estimator (86) with a section spacing of 5 *µ*m,

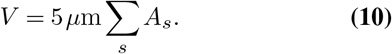

The 3D cross-K and cross-L functions were calculated using the 3D epithelial and T cell coordinate sets *E* and *T* as,

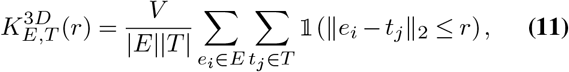

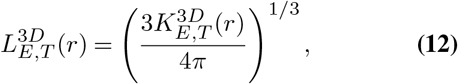

and evaluated over the same 0–20 *µ*m range, with its AUC obtained by trapezoidal integration. Neighbor counts in both dimensionalities were evaluated by KD-tree radius queries, and no edge correction was applied (Fig. 5b–h).

#### PDAC–IPMN immune microenvironment analysis

To compare local T cell organization relative to epithelial cells between IPMN and IPMN-associated PDAC, in individual 2D sections and in the corresponding 3D reconstructed volume, we selected an ROI that contained both lesions and spanned 11 serial sections. PDAC and IPMN compartments were manually annotated in each section and expanded outward by 100 *µ*m to define the surrounding lesion-associated microenvironments. Where the expanded regions overlapped, each pixel was assigned to the lesion whose original, unexpanded annotation was nearest, keeping the two microenvironments mutually exclusive. H&E-predicted cells were assigned to a microenvironment by their section-wise coordinates, followed by assembling of 3D cell maps from the registered serial-sections, and obtaining the 3D *x, y, z* cell coordinates obtained by spot detection in Imaris, using the settings described above.

Local cellular neighborhoods were represented by Delaunay triangulation (51) of the cell coordinates, constructed both within individual sections and across the 3D reconstructed volume, with edges longer than 50 *µ*m removed to restrict the graphs to local relationships. For T cell *i*, the interaction score with target class *c* is defined as the fraction of its retained neighbors belonging to that class,

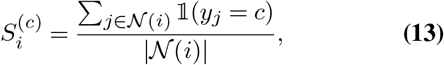

where *N*(*i*) denotes the retained graph neighbors of T cell *i* and *y*_*j*_ the predicted cell type *c* of neighbor *j*. T cells with no retained neighbors were assigned a score of zero. We report the T cell *→*Epithelial (T *→*E) and T cell*→*other scores, which quantify local association of T cells with epithelial and ‘other’ cells respectively.

Variation in T cell–epithelial organization across tissue depth was quantified from T*→*E scores calculated independently within each section. For each section and microenvironment, a 2D Delaunay graph was built from the section-specific coordinates with the same 50 *µ*m edge cutoff, and the T*→*E score summarized as the mean over all T cells in that microenvironment. To characterize the significance of the score, a random-mixing expectation was generated for each of these by permuting cell type labels 1,000 times while preserving the observed graph structure, cell positions, neighborhood sizes, and section-specific composition, recalculating the mean T*→*E score after each permutation. The expectation and its 95% interval, defined by the 2.5th and 97.5th percentiles, were summarized from the resulting null distribution, and observed section-wise scores compared against them across tissue depth for both microenvironments.

Heterogeneity in local T cell association across the reconstructed tissue in 3D was characterized from the distributions of individual T cell interaction scores, summarized separately for the PDAC- and IPMN-associated microenvironments, with mean scores reported as descriptive summaries (Fig. 5i–o).

#### Three-dimensional lymphoid aggregate detection, tracking, and quantification

Because individual lymphoid aggregates can extend across multiple serial sections, their three-dimensional extent was recovered by identifying section-wise lymphocyte clusters and linking corresponding profiles across tissue depth. This analysis used the same lymphocyte-rich, PDAC-containing ROI as the L-function analysis described above.

Candidate aggregates were identified independently within each section from H&E-predicted T and B cells. Their centroids were pooled and clustered using HDBSCAN (87, 88) with a minimum cluster size of 300 cells, minimum samples of 30, and excess-of-mass cluster selection. Because spatially distinct dense clusters can be connected by sparse lymphocyte bridges, such components were separated by a DBSCAN-based refinement (89) with a neighborhood radius of 18.75 *µ*m (75 pixels for a resolution of 0.25*µ*m/px) and a minimum of 10 samples. An *α*-shape boundary was then estimated around the dense core of each candidate cluster, with boundary estimation restricted to the denser portion of the cluster by local nearest-neighbor density so that sparsely distributed peripheral cells could not disproportionately extend it. All H&E-predicted T and B cells with centroids inside the resulting boundary were recovered as aggregate-associated lymphocytes. To focus the analysis on larger, dense aggregates, we retained only those containing at least 400 lymphocytes at a density of at least 3.5 cells per 1,000 *µ*m^2^.

Section-wise aggregate profiles were linked across tissue depth by boundary overlap between adjacent registered sections, and assigned to the same 3D track when their intersection over union reached 0.30. One-to-many matching was permitted so that track continuity was preserved where a single profile appeared as multiple spatial components in an adjacent section. Missing profiles across gaps of up to 10 sections were recovered under a relaxed lymphocyte-count threshold, and a further overlap-based merging step joined fragmented tracks while preserving distinct aggregate components occurring within the same section. Only tracks spanning at least three serial sections were retained.

For each retained track, section-wise boundaries were exported as binary masks for 3D visualization in Imaris. Cell composition was quantified from all classified cells with centroids inside the final boundaries across the sections comprising the track, including epithelial, T, B, and ‘other’cells, although detection itself used only the T and B cell populations. Cell-type-specific one-pixel centroid masks were assembled across sections and used for spot detection in Imaris using the settings described above to obtain 3D counts per track.

Aggregate proximity to PDAC was compared between individual sections and the 3D tissue reconstruction using LA2 as a representative aggregate, with epithelial cells inside the original PDAC annotations as the tumor reference population. For the section-specific analysis, the minimum in-plane Euclidean distance between LA2-associated lymphocytes and PDAC-associated epithelial cells was calculated independently within each section and the resulting values summarized by their median. For the 3D analysis, the minimum Euclidean distance between the same two populations was calculated across the reconstructed tissue in 3D from Imaris-derived *x, y, z* coordinates using KD-tree nearest-neighbor queries (Fig. 6c,d).

## Data availability

Publicly available datasets used in this study, together with their original sources and accession information, are listed in Supplementary Table 1.

## Code availability

The SpaReg source code and scripts used in this study will be made publicly available at https://github.com/uttamLab/SpaReg.

## Acknowledgements

We thank Yael Arbely for assistance with figure preparation.

## Author contributions

S.U. conceived the study. R.P., T.J. and S.U. developed the SpaReg and H&E-based cell classification method. R.P. and T.J. performed the analyses and generated the results. R.R. and E.B. contributed to data generation. S.W. contributed to 3D rendering and figure preparation. T.R.S. and A.S. provided tissue specimens and pathology expertise, and A.S. performed pathology annotations. S.U. supervised the study and acquired funding. R.P., T.J. and S.U. wrote the manuscript. All authors reviewed and approved the final manuscript.

---

**Supplementary Table 1.** Datasets used in this study. *µ*m/px denotes the native imaging resolution; for spatial-transcriptomics platforms, the unit of measurement is the spot or cell.

| Tissue / dataset | Modality / platform | Resolution | No. of sections | Size | Shown in |
| --- | --- | --- | --- | --- | --- |
| PDAC (cell classifier training and evaluation) | H&E, Leica Aperio AT2 (40×)<br>CyCIF, Nikon Ti2E (20×/0.75 NA) | 0.25 $\mu\text{m}/\text{px}$ (H&E)<br>0.32 $\mu\text{m}/\text{px}$ (IF) | 1 | 8 regions<br>181,976 nuclei | Fig. 1c–e<br>Fig. 4a–b |
| PDAC | H&E, Leica Aperio AT2 (40×) | 0.25 $\mu\text{m}/\text{px}$ | 320 | $\sim 2.77 \text{ cm}^2/\text{section}$ | Fig. 2a–b<br>Fig. 4c–d<br>Fig. 5<br>Fig. 6<br>Supplementary Video 4 |
| CRC | H&E, Leica Aperio AT2 (40×) | 0.25 $\mu\text{m}/\text{px}$ | 307 | $\sim 2.91 \text{ cm}^2/\text{section}$ | Fig. 2c–d<br>Extended Data Fig. 2 |
| HTAN CRC (hybrid) (3) | Interleaved H&E and CyCIF | 0.25 $\mu\text{m}/\text{px}$ (H&E)<br>0.65 $\mu\text{m}/\text{px}$ (IF) | 21 H&E<br>24 IF | $\sim 1.126 \text{ cm}^2/\text{section}$ | Fig. 2e–f<br>Supplementary Video 3 |
| Cross-platform mouse brain (46–48) | CosMx, Xenium 5K, Xenium, STARmap+, MERFISH, Visium HD, Visium | Platform-dependent | 7 | – | Fig. 2g |
| Zhuang-ABCA-1 (90) | MERFISH, whole mouse brain | Single cells | 129 | $\sim 2.64$ million cells<br>1,122 genes | Fig. 2h<br>Extended Data Fig. 4 |
| Murine prostate benchmark (16) | H&E | 0.46 $\mu\text{m}/\text{px}$ | 260 | $\sim 0.23 \text{ cm}^2/\text{section}$ | Fig. 3a–b<br>Extended Data Fig. 5<br>Extended Data Fig. 6 |
| Human DLPFC (91)<br>Donor III, sections 151673–151676 | 10x Visium | Spots | 4 | $\sim 3,561$ spots/section<br>33,538 genes | Fig. 3c<br>Extended Data Fig. 7 |
| Mouse MHPR (92) | MERFISH | Single cells | 5 | $\sim 5,663$ cells/section<br>155 genes | Fig. 3d |
| High-grade serous ovarian carcinoma (HGSOC) | H&E, Leica Aperio AT2 (40×) | 0.25 $\mu\text{m}/\text{px}$ | 220 | $\sim 3.19 \text{ cm}^2/\text{section}$ | Extended Data Fig. 3a<br>Supplementary Video 2 |
| Benign fallopian tube tissue | H&E, Leica Aperio AT2 (40×) | 0.25 $\mu\text{m}/\text{px}$ | 200 | $\sim 4.87 \text{ cm}^2/\text{section}$ | Extended Data Fig. 3b |
| Cholangiocarcinoma | H&E, Leica Aperio AT2 (40×) | 0.25 $\mu\text{m}/\text{px}$ | 80 | $\sim 5.3 \text{ cm}^2/\text{section}$ | Supplementary Video 1 |

